# tRNAnalysis: A flexible pre-processing and next-generation sequencing data analysis pipeline for transfer RNA

**DOI:** 10.1101/655829

**Authors:** Anna James-Bott, Adam P. Cribbs

## Abstract

Many tools have been developed to analyse small RNA sequencing data, however it remains a challenging task to accurately process reads aligning to small RNA due to their short-read length. Most pipelines have been developed with miRNA analysis in mind and there are currently very few workflows focused on the analysis of transfer RNAs. Moreover, these workflows suffer from being low throughput, difficult to install and lack sufficient visualisation to make the output interpretable. To address these issues, we have built a comprehensive and customisable small RNA-seq data analysis pipeline, with emphasis on the analysis of tRNAs. The pipeline takes as an input a fastq file of small RNA sequencing reads and performs successive steps of mapping and alignment to transposable elements, gene transcripts, miRNAs, snRNAs, rRNA and tRNAs. Subsequent steps are then performed to generate summary statistics on reads of tRNA origin, which are then visualised in a html report. Unlike other low-throughput analysis tools currently available, our high-throughput method allows for the simultaneous analysis of multiple samples and scales with the number of input files. tRNAnalysis is command line runnable and is implemented predominantly using Python and R. The source code is available at https://github.com/Acribbs/tRNAnalysis.

## Introduction

Transfer RNAs (tRNAs) are non-coding RNAs that implement the genetic code by delivering amino acids to ribosomes during translation. Most genomes contain distinct tRNAs for all 61 codons and these are encoded across multiple sites throughout the genome. Sequencing of tRNA is challenging both experimentally and computationally. The main experimental challenges arise from a stable secondary and tertiary structure, making library preparation difficult [1]. Therefore, efficient library preparation methods must be employed to overcome the ridged structure of tRNA that usually limit the use of standard library prep methods for sequencing [2]. With respect to data analysis, the challenges come from overcoming the reverse transcription errors introduced by chemical modifications and accurately mapping reads to tRNA genomic regions, given their multiple identical and almost identical genomic loci [3]. Typically, most mapping strategies for gene expression analysis only report read alignments with unique best matches and thus discard reads mapping to tRNA altogether. As a consequence, specialist mapping strategies to accurately map tRNAs have been proposed [4, 5]. Specifically, Hoffmann *et al* (2018) have recently proposed a two pass mapping strategy that first maps reads to a tRNA masked genome then secondly these unmapped reads are aligned directly to merged tRNA clusters [3].

While computational pipelines have been developed in the past for small RNA-seq data analysis [6], there is currently a limited number of small RNA-seq pipelines focusing on tRNA data analysis. While there have been low-throughput implementations to aid tRNA analysis, such as tDRmapper [4], tRF2Cancer [7], SPORTS1.0 [8] and MINTmap [5], there is now a significant unmet requirement for high-throughput approaches to analyse tRNA data. Firstly, it is desirable to have a single pipeline with the flexibility to perform read quality control, mapping to both general genomic and tRNA features, in addition to performing qualitative and quantitative analyses on tRNAs within a sample. Secondly, with falling sequencing costs, it is now common for small RNA-seq libraries to consist of many biological and technical repeats and be sequenced at a much greater depth. Thirdly, it is desirable to have a pipeline that generates an appropriate level of detailed reporting output, which includes appropriate visualisation and publication quality figures.

tRNAnalysis is a pipeline written using the CGAT-core workflow manager [9]. It is seamlessly integrated and runs from a single launch command, while also having the modularity of being able to run any individual task within the pipeline. tRNAnalysis implements best practice mapping strategies to allow accurate mapping of tRNA reads [3]. The pipeline is optimised so that it can process all input fastq files in parallel, produce detailed logging information, and can be ran locally or using a high-performance cluster. tRNAnalysis can be installed using Conda and a Docker image is also provided with all of the software and packages installed. Users can therefore plug and play the pipeline without having to install numerous dependencies. Finally, we provide a user-friendly html report to visualise qualitative and quantitative outputs from our pipeline.

## Methods

tRNAnalysis automates and integrates best-practice small RNA sequencing analysis, allowing for automatic cluster submission and parallelisation. The pipeline unifies standard software for its functions, such that the pipeline provides appropriate default settings, with the option to customise pipeline configuration parameters and job resources as required. The workflow is written predominantly in python and R, using CGAT-ruffus decorators [10] and CGAT-core [9] as the workflow management system. The pipeline runs via a single command line interface, and the main steps in our analysis being:

- Read pre-processing and quality assessment
- Mapping of reads
- tRNA quantitative and qualitative analysis
- Downstream analyses and report generation

### Read pre-processing and quality control

tRNAnalysis accepts raw sequencing data (single-end fastq files) as input and integrates several tools for read quality checking and filtering, including CGAT tools [11], FastQC [12] and FastQ Screen [13] and applies quality control and pre-processing steps to each file. Given the short length of small RNAs captured during library preparation, Trimmomatic [14] is used to remove adapter and filter reads that fall short of quality thresholds. Results for each sample are then summarised with MultiQC [15].

### Mapping of reads

The input for mapping is a collection of pre-processed fastq files and a list of gencode annotations, which are supplemented with automatically downloaded RNA repeats (including RNA, tRNA, rRNA, snRNA, srpRNA) from the UCSC database [16]. Firstly, mapping is performed against the genome using Bowtie [17] to obtain a global representation of RNA types. For effective tRNA mapping, tRNAnalysis implements the best-practice mapping strategy first proposed by Hoffmann *et al* (2018), in which tRNA loci are masked from the genome and instead, intron-less tRNA precursor sequences are appended as artificial chromosomes [3]. In first-pass mapping, reads that overlap the boundaries of mature tRNAs are extracted. In a subsequent round of mapping, the remaining reads are mapped to a tRNA-masked target genome that is augmented by representative mature tRNA sequences.

### tRNA qualitative and quantitative analysis

We provide RNA profile analysis that summarises the output of the read alignments mapping to various RNA sequences (e.g. miRNA, piRNA, snoRNA, lncRNA e.c.t.) by counting reads to features with featureCounts [18]. Plots are then generated for each sample, where the positional coverage counts are plotted relative to the exon, upstream and downstream regions of the tRNAs. These plots can then be used to infer the levels of tRNA fragments within a sample. Using the nomenclature first proposed by Selitsky *et al* (2015), if the primary tRNA sequence is < 41 nts and >= 28 nts, then it is defined as a tRNA-half, while if it is > 14 nts and < 28 nts then it is defined as a tRNA-fragment [19]. We calculate the frequency of read end site relative to the tRNA length and plot these as bar graphs for each tRNA cluster type. For quantitative measurement of tRNA differences between groups of samples, we implement DESeq2 to perform differential expression analysis [20]. Finally, given there can be large sequence variations between tRNAs from different tissues as a consequence of RNA modifications [3, 21, 22], we also determine nucleotide misincorporations in the mapped reads. In order to accurately distinguish sequencing errors for true mismatches, we employ *samtools mpileup* to collate summary information in the mapped bam file and then compute likelihoods of misincorportation. This information is stored in a bcf file that is then parsed by *bcftools call*, which performs variant calling for each tRNA sequence [23]. The output is then stored as a vcf file and normalised for indels, then filtered for sequencing depth.

### Visualisation

In order to visualise the output of the pipeline, summary statistics are generated and then reports of these summaries are rendered in html format for visualisation using the Rmarkdown framework. Publication ready reports include figures such as coverage plots, volcano plots of differential expressed tRNAs, tables detailing tRNA modifications and bar graphs of tRNA frequencies. Once these standard reports have been generated, exploratory data analysis can be performed by modifying the Rmarkdown code, allowing for bespoke analysis, tweaking some parameters or to generate more specific publication-quality images.

## Implementation

tRNAnalysis is implemented in Python 3 and installable via Conda and Pypi. We have successfully deployed and tested the code on OSX, Red Hat and Ubuntu. The pipeline requires a functional version of Bowtie for mapping of reads to a specified reference genome and tRNA sequences. Compression of SAM files to BAM format, in addition to variant calling using *mpileup*, requires SAMtools [23]. The pipeline is constructed using the CGAT-core workflow engine (tested using v0.5.10, [9]), with python decorators from CGAT-ruffus (tested using v2.8.1, [10]) being used to track the input and output files. For full functionality, tRNAnalysis requires the statistical programming language R (tested using v3.4.1 and v3.5.1). However, all software and dependencies required to run tRNAnalysis can be installed using the bioconda framework [24]. Moreover, we also have tRNAnalysis as a docker container, allowing seamless plug and play without pre-installing the numerous dependencies.

We have made tRNAnalysis and associated repositories open source under the MIT licence, allowing full and free use for both commercial and non-commercial purposes. Our software is fully documented (https://pypi.org), version controlled and tested using continuous integration (https://travis-ci.com/Acribbs/single-cell). We welcome community participation in code development and issue reporting through GitHub.

## Results

To allow users to familiarise themselves with the functionality of tRNAnalysis, we have included a tutorial example that can be accessed in the github documentation. However, in order to validate the accuracy of our methods we have used our pipeline to analyse previously generated datasets from Chiou *et al* (2018) [25]. This dataset comprises small RNA samples isolated from activated T cells and their associated extracellular vesicles. Using our workflow, we were able to confirm the main findings of Chiou *et al* (2018), mainly that specific tRNA fragments are enriched in extracellular vesicles and released by activated T cells (Figure 1). Furthermore, using our workflow we were able to present a more detailed evaluation of the different tRNA types, mapping reads collapsed across all tRNAs (Figure 1A), by codon (not shown) and by amino acid (Figure 1B). Our pipeline also allows the user to quickly evaluate the relative proportion of tRNA fragments (Figure 1C) and tRNA halves (Figure 1D) in a sample. Furthermore, we also demonstrated that tRNAnalysis can be used to accurately quantify tRNA differential expression between cells and extracellular vesicles (Figure 2).

**Figure 1:**
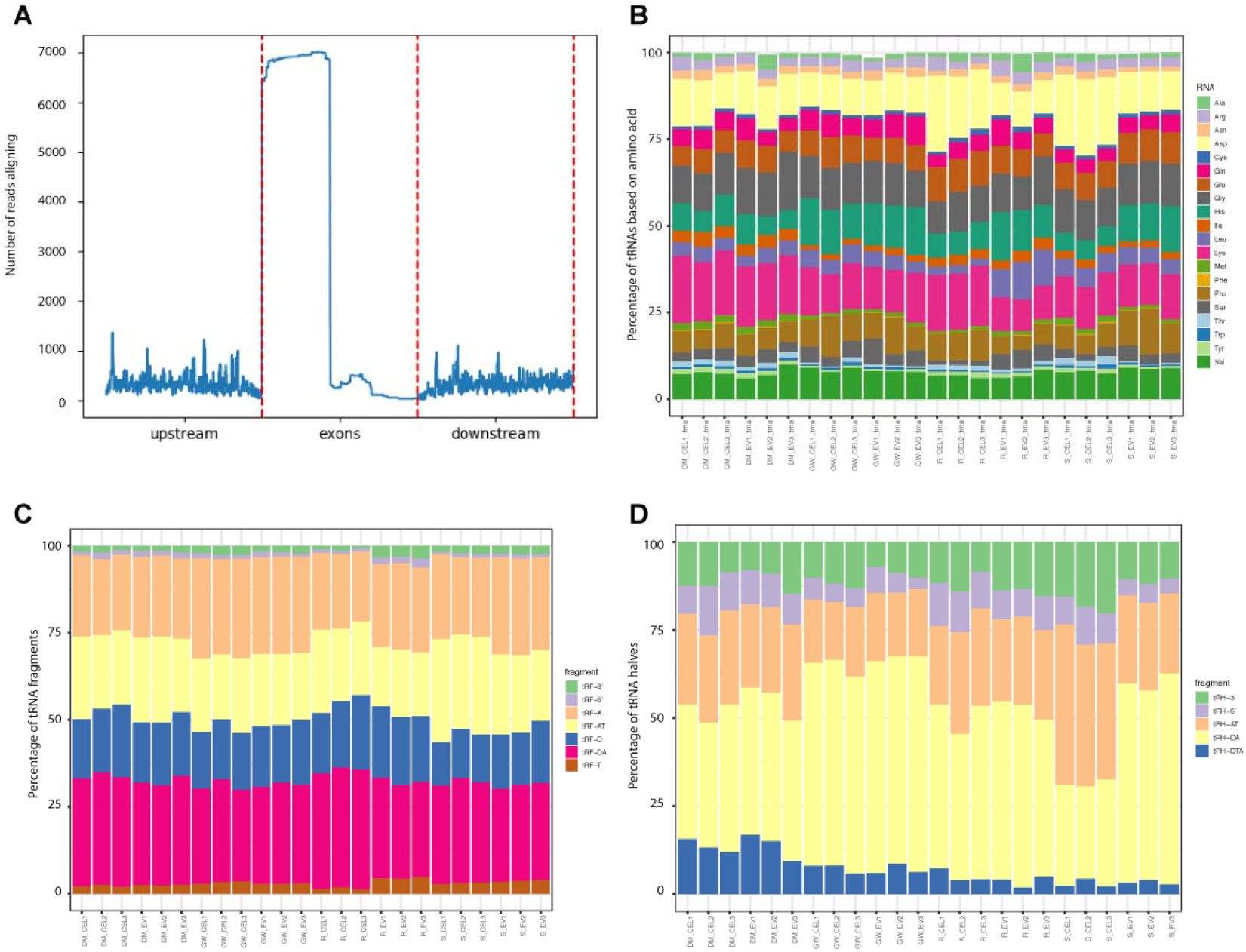
Example of qualitative plots generated by tRNAnalysis of reads mapping to tRNA. A: Gene profile figures showing read counts of tRNA sequences from 5’ (upstream) to 3’ (downstream) of all collapsed tRNAs. B: Proportion of sequencing reads mapped to different small RNA samples in the Chiou *et al (2018)* data and collapsed by amino acid. C: Proportion of sequencing reads mapped to different tRNA fragments. D: Proportion of sequencing reads mapped to different tRNA halves.

**Figure 2:**
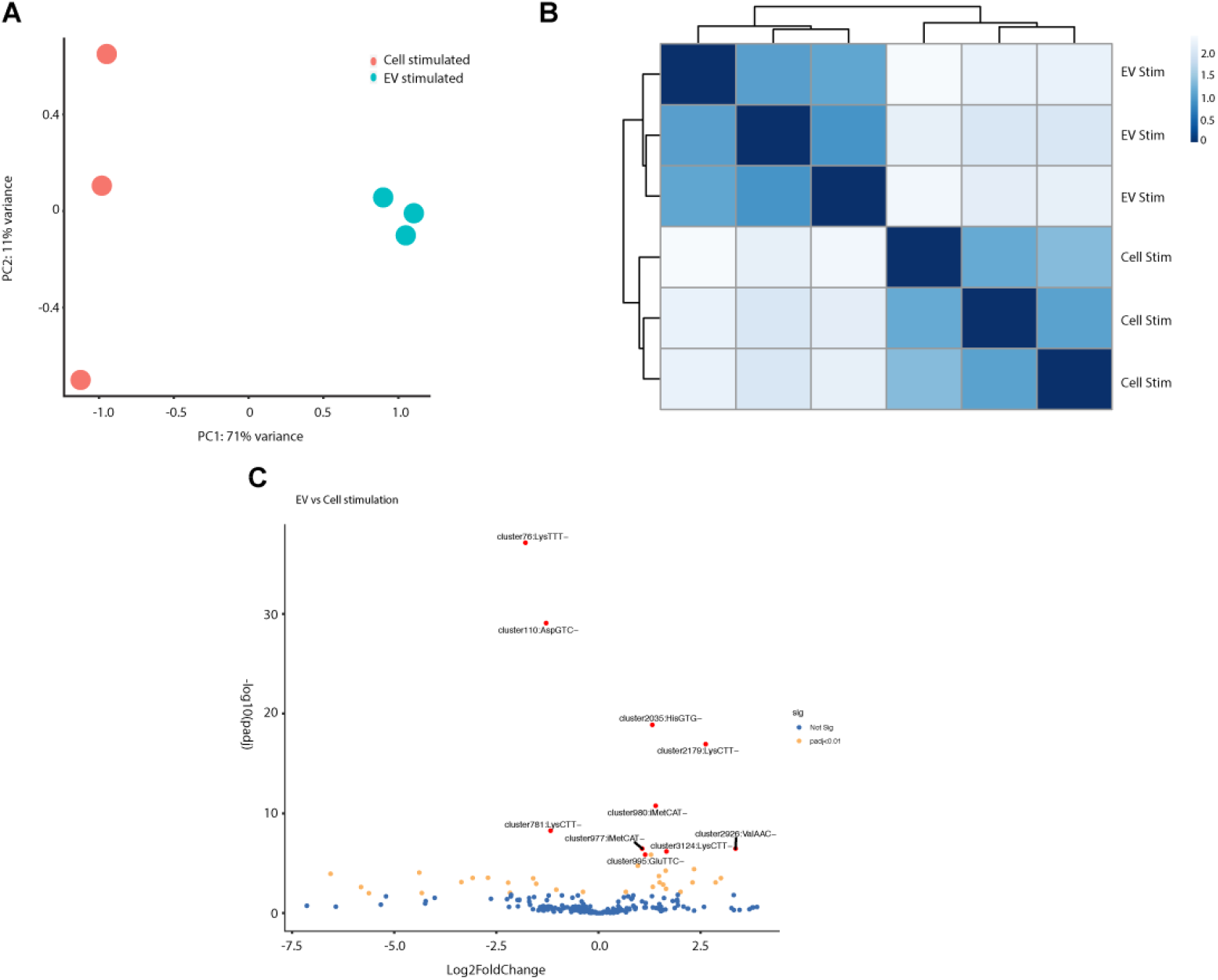
Example of quantitative plots of the differential expression analysis generated by tRNAnalysis. A: Principle component analysis demonstrating clear separation between stimulated cells and extracellular vesicles. B: Sample-sample distances displayed as a heatmap. The heatmap was generated by DESeq2 software package and shows the Euclidean distance between each sample. C: Volcano plot showing relative up- and down-regulation of tRNA species in MLEV when compared to SEV

## Discussion

We have developed a one-stop pipeline for the processing and analysis of small RNA-seq data, with a particular emphasis on the analysis of reads aligning to tRNAs. tRNAnalysis is expansible and robust, and can be run locally or scaled up and ran on an HPC cluster. Additionally, the pipeline is portable since it is distributed as a conda package and a docker image. Currently, tRNAnalsyis supports data generated from all major RNA-seq platforms and implements best practice accurate mapping of tRNA reads [3]. The pipeline encourages a best practice reproducible research approach, by implementing exploratory data analysis using the R statistical programming framework, which is initially generated by the pipeline. Wrapping of prototyped analysis Rmarkdown files allows the rapid reproduction of analyses and reuse of code for multiple tRNA datasets. To illustrate a simple use case of tRNAnalysis, we have included an example small RNA-seq dataset and tutorial, which can be accessed from within the github repository (https://github.com/Acribbs/tRNAnalysis). tRNAnalysis is currently being used to process data in numerous ongoing projects and is under active development to meet the demands of the burgeoning small RNA sequencing field.

## Data availability

All data underlying the results are available as part of the article and no additional source data are required

## Software availability

Source code is available from: https://github.com/Acribbs/tRNAnalysis

Archived source code at time of publication: https://zenodo.org/record/3234845#.XO60ti2ZPPA

Licence: MIT Licence

## Funding

APC was supported through funding from a CRUK Oxford Centre Development Fund award (CRUKDF-0318-AC(AZ)) and as a trainee in the LEAN program grant from the Leducq Foundation.

## References

1. Kim, S.H., et al., Three-dimensional structure of yeast phenylalanine transfer RNA: folding of the polynucleotide chain. Science, 1973. 179(4070): p. 285–8.

2. Shigematsu, M., et al., YAMAT-seq: an efficient method for high-throughput sequencing of mature transfer RNAs. Nucleic Acids Res, 2017. 45(9): p. e70.

3. Hoffmann, A., et al., Accurate mapping of tRNA reads. Bioinformatics, 2018. 34(13): p. 2339.

4. Selitsky, S.R. and P. Sethupathy, tDRmapper: challenges and solutions to mapping, naming, and quantifying tRNA-derived RNAs from human small RNA-sequencing data. BMC Bioinformatics, 2015. 16: p. 354.

5. Loher, P., A.G. Telonis, and I. Rigoutsos, MINTmap: fast and exhaustive profiling of nuclear and mitochondrial tRNA fragments from short RNA-seq data. Sci Rep, 2017. 7: p. 41184.

6. Wu, X., et al., sRNAnalyzer-a flexible and customizable small RNA sequencing data analysis pipeline. Nucleic Acids Res, 2017. 45(21): p. 12140–12151.

7. Zheng, L.L., et al., tRF2Cancer: A web server to detect tRNA-derived small RNA fragments (tRFs) and their expression in multiple cancers. Nucleic Acids Res, 2016. 44(W1): p. W185–93.

8. Shi, J., et al., SPORTS1.0: A Tool for Annotating and Profiling Non-coding RNAs Optimized for rRNA- and tRNA-derived Small RNAs. Genomics Proteomics Bioinformatics, 2018. 16(2): p. 144–151.

9. Cribbs, A., et al., CGAT-core: a python framework for building scalable, reproducible computational biology workflows [version 1; peer review: 1 approved, 1 approved with reservations]. F1000Research, 2019. 8(377).

10. Goodstadt, L., Ruffus: a lightweight Python library for computational pipelines. Bioinformatics, 2010. 26(21): p. 2778–9.

11. Sims, D., et al., CGAT: computational genomics analysis toolkit. Bioinformatics, 2014. 30(9): p. 1290–1.

12. Andrews, S., FastQC: a quality control tool for high throughput sequence data. 2010.

13. Wingett, S.W. and S. Andrews, FastQ Screen: A tool for multi-genome mapping and quality control. F1000Res, 2018. 7: p. 1338.

14. Bolger, A.M., M. Lohse, and B. Usadel, Trimmomatic: a flexible trimmer for Illumina sequence data. Bioinformatics, 2014. 30(15): p. 2114–20.

15. Ewels, P., et al., MultiQC: summarize analysis results for multiple tools and samples in a single report. Bioinformatics, 2016. 32(19): p. 3047–8.

16. Karolchik, D., et al., The UCSC Table Browser data retrieval tool. Nucleic Acids Res, 2004. 32(Database issue): p. D493–6.

17. Langmead, B., et al., Ultrafast and memory-efficient alignment of short DNA sequences to the human genome. Genome Biol, 2009. 10(3): p. R25.

18. Liao, Y., G.K. Smyth, and W. Shi, featureCounts: an efficient general purpose program for assigning sequence reads to genomic features. Bioinformatics, 2014. 30(7): p. 923–30.

19. Selitsky, S.R., et al., Small tRNA-derived RNAs are increased and more abundant than microRNAs in chronic hepatitis B and C. Sci Rep, 2015. 5: p. 7675.

20. Love, M.I., W. Huber, and S. Anders, Moderated estimation of fold change and dispersion for RNA-seq data with DESeq2. Genome Biol, 2014. 15(12): p. 550.

21. Vilmi, T., et al., Sequence variation in the tRNA genes of human mitochondrial DNA. J Mol Evol, 2005. 60(5): p. 587–97.

22. Lin, B.Y., P.P. Chan, and T.M. Lowe, tRNAviz: explore and visualize tRNA sequence features. Nucleic Acids Res, 2019.

23. Li, H., et al., The Sequence Alignment/Map format and SAMtools. Bioinformatics, 2009. 25(16): p. 2078–9.

24. Gruning, B., et al., Bioconda: sustainable and comprehensive software distribution for the life sciences. Nat Methods, 2018. 15(7): p. 475–476.

25. Chiou, N.T., R. Kageyama, and K.M. Ansel, Selective Export into Extracellular Vesicles and Function of tRNA Fragments during T Cell Activation. Cell Rep, 2018. 25(12): p. 3356–3370 e4.

